# Alteration of perivascular spaces in early cognitive decline

**DOI:** 10.1101/2020.01.30.927350

**Authors:** Farshid Sepehrband, Giuseppe Barisano, Nasim Sheikh-Bahaei, Jeiran Choupan, Ryan P Cabeen, Malcolm S Crawford, Wendy J Mack, Helena C Chui, John M Ringman, Arthur W Toga, for the Alzheimer’s Disease Neuroimaging Initiative

## Abstract

Vascular contributions to early cognitive decline are increasingly recognized, prompting further investigation into the nature of related changes in perivascular space. Using magnetic resonance imaging, we show that, compared to a cognitively normal sample, individuals with early cognitive dysfunction have altered perivascular space presence and distribution, irrespective of Amyloid-β. Surprisingly, we noted decreased perivascular space presence in the anterosuperior medial temporal lobe, which was associated with neurofibrillary tau tangle deposition in the entorhinal cortex, one of the hallmarks of early Alzheimer’s disease pathology. Our results suggest that anatomically-specific alteration of the perivascular spaces may provide an early biomarker of cognitive impairment in aging adults.

## Introduction

One of the potential pathophysiological hypotheses for cognitive decline is represented by neurovascular dysfunction. Preclinical evidence in a pericyte-deficient mouse model supports the hypothesis that blood-brain barrier (BBB) breakdown leads to extravascular accumulation of blood-derived toxic fibrin deposits, with enlargement of perivascular spaces (PVS) and early white matter damage at the time when hypoxic changes are undetectable in the cortex [1]. Similarly, a recent study in humans showed that individuals with early cognitive dysfunction develop brain capillary and BBB damage in the hippocampus, suggesting that BBB breakdown is an early biomarker of cognitive decline [2]. In addition, previous studies in animals suggest that the glymphatic (glia-lymphatic) system, whose main pathway is represented by the perivascular spaces, is a substantial factor in the net clearance of Aβ [3, 4]; reduction of efflux of interstitial fluid to the cerebrospinal fluid (CSF) in PVS could lessen Aβ clearance [5–7]. Such an impaired clearance from the brain parenchyma could in part explain the accumulation of Aβ [5]. Aβ deposition in the wall of leptomeningeal and intraparenchymal cerebral arteries is a common observation in Alzheimer’ disease (AD), that is thought to be associated with the enlargement of PVS as routes of Aβ clearance [8–10]. PVS changes have been linked to vascular diseases, including cerebral small vessel disease, cerebral amyloid angiopathy, BBB breakdown, hypertension and lacunar stroke [1,11–17]. Furthermore, an emerging body of evidence suggests vascular changes are linked to both clearance system dysfunction [18–23] and vasculotoxic effects of Aβ and tau [10, 24].

Magnetic resonance imaging (MRI) studies have found significantly larger PVS in AD patients compared with controls in the centrum semi-ovale of the white matter (CSO-WM), but not in the basal ganglia [25, 26]. Interestingly, Banerjee et al. found that the perivascular spaces in the white matter were associated with AD independently of the amyloid burden [26]. These findings are limited to clinical scoring of PVS, which is based on counting total number of observed PVS in the white matter. The anatomical distribution and morphological features of PVS alterations in white matter however remain largely unexplored. To date, it has not been demonstrated whether PVS may represent an early biomarker of cognitive decline, nor have relationships of PVS with other pathological markers of AD, including Aβ and tau been reported.

To examine whether and to what extent PVS is altered in early cognitive decline, we performed a comprehensive analysis of the multi-modal data of the Alzheimer’s Disease Neuroimaging Initiative 3 (ADNI-3) participants (N=596). Since Aβ and tau can both lead to cerebral blood vessel abnormalities [27–29], we mapped PVS using MRI data and analyzed the relationship between PVS volume fraction and Aβ and tau accumulation as measured in positron emission tomography (PET) images. Additionally, we investigated whether PVS distribution and appearance on MRI are influenced by the *APOE* gene, whose ε4 variant represents one of the strongest genetic risk factors for AD [30].

## Method

### Study Participants

Data used in the preparation of this article were obtained from the Alzheimer’s Disease Neuroimaging Initiative 3 (ADNI-3) database (http://adni.loni.usc.edu). Data from ADNI-3 cohort [32] was used, in which both T1-weighted (T1w) and fluid-attenuated inversion recovery (FLAIR) structural magnetic resonance image (MRI) were available. T1w and FLAIR images for 685 participants were downloaded from the ADNI database (http://adni.loni.usc.edu) [33]. Demographic information, clinical and cognitive assessment, positron emission tomography (PET), cardiovascular-related health measures and cerebrospinal fluid (CSF) biomarkers were collected. 51 participants who met the AD categorization criteria were excluded (as described in *Clinical and cognitive assessment* section). An additional 37 participants who failed MRI quality control were excluded (as described in *Mapping perivascular spaces* section), resulting in a total of 596 participants.

### Inclusion/Exclusion Criteria

All participants must not have had any significant neurologic disease other than suspected incipient Alzheimer’s disease; excluded neurologic condition included Parkinson’s disease, multi-infarct dementia, Huntington’s disease, normal pressure hydrocephalus, brain tumor, progressive supranuclear palsy, seizure disorder, subdural hematoma, multiple sclerosis, or history of significant head trauma followed by persistent neurologic deficits or known structural brain abnormalities. Additional and detailed inclusion and exclusion criterion are described in the ADNI-3 protocol [32] and also copied in **Supplementary File 1**.

### Clinical and cognitive assessments

The ADNI clinical dataset comprises clinical information about each participant including recruitment, demographics, physical examinations, and cognitive assessment data. Demographic information including age, sex and the years of education were gathered, which were used as covariates in all statistical analysis. A clinical interview and a general physical exam were conducted. In addition, respiration rate, blood glucose level (after overnight fasting), and systolic and diastolic pressures were acquired in the 30-day interval of the imaging visit. The cognitive assessment was performed in accordance with published standardization procedures, including standardized interview and assessment with the participant and a knowledgeable informant.

#### Clinical dementia rating (CDR)

Six categories of cognitive functioning (memory, orientation, judgment and problem solving, community affairs, home and hobbies, and personal care) were assessed, in which the CDR describes five degrees of impairment [34, 35]. Participant CDR scores were obtained through a standardized interview and assessment with the participant and a knowledgeable informant.

#### Mini-Mental State Examination (MMSE)

The MMSE scale evaluates orientation, memory, attention, concentration, naming, repetition, comprehension, and ability to create a sentence and to copy two overlapping pentagons [36]. Participant MMSE scores were obtained through standardized interview and assessment with the participant and a knowledgeable informant Participant categorization is detailed in the ADNI-3 protocol document. In brief:

#### Cognitively normal (CN)

Participants were considered CN if they had normal memory function, which was assessed using education-adjusted cutoffs on the Logical Memory II subscale from the Wechsler Memory Scale [37]. Participants must also have an MMSE score between 24 and 30 and a CDR of 0 (memory box score must be 0). CN participants were considered cognitively normal based on an absence of significant impairment in cognitive functions or daily living activities.

#### Mild cognitive impairment (MCI)

Participants were classified as MCI if they expressed subjective memory concern, with a score below the education-adjusted cutoffs on the Logical Memory II subscale from the Wechsler Memory Scale. The same MMSE score range as CN was used for the MCI group. Participants must have a global CDR of 0.5 and the memory box score must be at least 0.5. General cognition and functional performance of the MCI participants must be sufficiently preserved such that clinical diagnosis of Alzheimer’s disease could not be made.

#### Alzheimer’s disease (AD)

Participants categorized as AD were excluded from this study, to ensure the focus on early cognitive decline. In brief, participants were considered AD if they scored below the education-adjusted cutoff of the Wechsler Memory Scale, had an MMSE score between 20 to 24 and global CDR>= 0.5 [32].

### Genotyping

A 7 mL sample of blood was taken in ethylenediaminetetraacetic acid (EDTA)-containing vacutainer tubes and genomic DNA was extracted at Cogenics (now Beckman Coulter Genomics) using the QIAamp DNA Blood Maxi Kit (Qiagen, Inc, Valencia, CA) following the manufacturer’s protocol. All DNA extraction and genotyping was done blinded to group assignment. The two SNPs (rs429358, rs7412) that define the ε2, ε3, and ε4 alleles of the apolipoprotein E (*APOE*) were genotyped by PCR amplification and HhaI restriction enzyme digestion. The genotype was resolved on 4% Metaphor Gel and visualized by ethidium bromide staining.

### Positron emission tomography

Amyloid PET data was used to categorize participants to Aβ+ and Aβ-groups. Amyloid PET analysis was performed according to UC Berkeley PET methodology for quantitative measurement [38–41]. Participants were imaged by Florbetapir (18 F-AV-45, Avid), or 18 F-Florbetaben (NeuraCeq, Piramal). All PET images were co-registered on T1w MRI to align brain parcellation boundaries. Quantitative measurement was done based on Standard Uptake Value ratio (SUVR). The tracer uptake in the main four cortical regions: frontal, anterior/posterior cingulate, lateral parietal, lateral temporal and additional five reference regions (cerebellar grey matter, whole cerebellum, brainstem/pons, eroded subcortical white matter, and a composite reference region) was measured. The whole cortical SUVR was then calculated by averaging the mean of tracer uptake across the 4 cortical regions and dividing by one of the reference regions.

Tau PET was also collected by Flortaucipir (_18_ F-AV-1451). Similar processing to Amyloid PET was performed to measure Tau PET SUVR. ADNI-3 Tau PET includes a broad set of regional SUVRs, which includes cortical and subcortical regions and eroded hemispheric WM, which enabled directed association analysis between perivascular space measures and Tau uptake. The weighted average SUVR values across Braak stage [42] regions were also used. Braak stage regions are listed in **Supplementary File 2**.

### Cerebral spinal fluid biomarkers

CSF Aβ_40_, CSF Aβ_42_ and Tau, phosphorylated Tau181 (p-Tau) measurement were completed using the Roche Elecsys cobase 601 fully automated immunoassay platform at the ADNI biomarker core (University of Pennsylvania). CSF biomarker samples were acquired in the 30-days interval of the imaging visit. All samples were aliquoted and stored at -80°C until assay. A 2D-ultra performance liquid chromatography tandem mass spectroscopy (UPLC/MS -MS) platform was used [43, 44], which was calibrated with a surrogate calibrator matrix prepared from artificial CSF plus 4 mg/mL bovine serum albumin.

### Magnetic resonance imaging

MRI imaging of the ADNI-3 was done exclusively on 3T scanners (Siemens, Philips, and GE) using a standardized protocol. 3D T1w with 1 mm3 resolution was acquired using an MPRAGE sequence (on Siemens and Philips scanners) and FSPGR (on GE scanners). For FLAIR images, a 3D sequence with similar resolution as T1w images was acquired, which provided the opportunity for accurate intrasubject intermodal co-registration. MPRAGE T1w MRI scans were acquired using the following parameters: TR = 2300 ms, TE = 2.98 ms, FOV = 240 × 256 mm2, matrix = 240 × 256 (variable slice number), TI = 900 ms, flip angle = 9, effective voxel resolution = 1 × 1 × 1 mm3. The FSPGR sequence was acquired using: sagittal slices, TR = 7.3 ms, TE = 3.01 ms, FOV = 256 × 256 mm2, matrix = 256 × 256 (variable slice number), TI = 400 ms, flip angle = 11, effective voxel resolution = 1 × 1 × 1 mm3. 3D FLAIR images were acquired using: sagittal slices, TR = 4,800 ms, TE = 441 ms, FOV = 256 × 256 mm2, matrix = 256 × 256 (variable slice number), TI = 1650 ms, flip angle = 120, effective voxel resolution = 1 × 1 × 1.2 mm3.

### MRI preprocessing and brain parcellation

T1w preprocessing and parcellation was done using the FreeSurfer (v5.3.0) software package, which is freely available [45], and data processing using the Laboratory of Neuro Imaging (LONI) pipeline system (http://pipeline.loni.usc.edu) [46–49], similar to [50, 51]. Brain volume and white matter mask were derived from the Desikan-Killiany atlas [52]. The parcellation was performed as part of the *recon-all* module of the FreeSurfer, which uses an atlas-based parcellation approach. Prior to parcellation, *recon-all* applies the following pre-processing steps: motion correction, non-uniform intensity normalization, Talairach transform computation, intensity normalization and skull stripping [52,53,62–65,54–61].

FLAIR images of each participants were corrected for non-uniform field inhomogeneity using N4ITK module [66] of Advanced Normalization Tools (ANTs) [67]. FLAIR images were then co-registered to T1w images using *antsIntermodalityIntrasubject* ANTs module.

### Mapping perivascular spaces

Perivascular spaces (PVS) were mapped from T1w images and then white matter hyperintensities, which were falsely segmented as PVS, were excluded using FLAIR images. PVS segmentation was done using an automated and highly reliable quantification technique that we developed and validated [68]. In brief, after preprocessing the data, T1w images were filtered using an adaptive non-local mean filtering technique [69]. The non-local mean technique measures the image intensity similarities by considering the neighboring voxels in a blockwise fashion, where the filtered image is 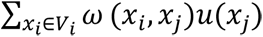. For each voxel (*x_j_*) the weight (ω) is measured using the Euclidean distance between 3D patches. The adaptive non-local mean filtering technique adds a regularization term to the above formulation to remove bias intensity of the Rician noise observed in MRI. Therefore the expected Euclidian distance between two noisy patches *N_i_* and *N_j_* is defined as: 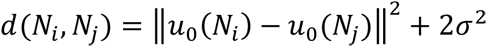, where 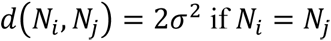. The Rician noise of the MRI images, calculated using robust noise estimation technique presented by Wiest-Daessle et al. [70], was used as the noise level for non-local filtering [69]. To preserve PVS voxels while removing the noise, filtering was applied only on high frequency spatial noises. This was achieved by using a filtering patch with a radius of 1 voxel, which removes the noise at a single-voxel level and preserves signal intensities that are spatially repeated [69].

Subsequently, we applied a Frangi filter [71] to T1w using the Quantitative Imaging Toolkit [72]. The Frangi filter estimates a vesselness measure for each voxel *V*(*s*) from eigenvectors λ of the Hessian matrix ℋ of the image:

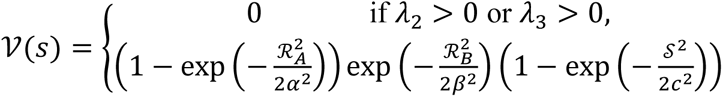

Where, 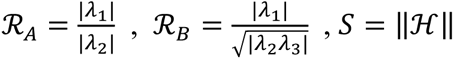.

Default parameters of α = 0.5, *β* = 0.5 and *c* were used, as recommended in [71]. The parameter *c* was set to half the value of the maximum Hessian norm. Frangi filter estimated vesselness measures at different scales and provided the maximum likeliness. The scale was set to a large range of 0.1 to 5 voxels in order to maximize the vessel inclusion. The output of this step is a quantitative map of vesselness in regions of interest and is taken to be the maximum across scales, as suggested in the original paper [71]. The range corresponds to specific levels in scale space that are searched for tubular structure feature detector. Thus, the outputs across voxels comprise vesselness measured across a range of filter scales. Then the optimum threshold of 1.5 [68] was used to segment the PVS voxels. For each segmented voxel, corresponding FLAIR voxels was checked, and if the FLAIR voxel value was a white matter hyperintensity, then the voxel was excluded from the final PVS mask. **Supplementary Figure 1** shows an example of the PVS map of one of the study participants.

T1w images were registered to the ANTs template nonlinear registration module [67, 73], and the same transformation matrix was applied on PVS masks. Then the normative PVS was generated for each of the CN and MCI groups (**Figure 1**). Two ROIs were then drawn on the template to enable statistical analysis of the PVS presence. The first ROI was drawn in the anterosuperior medial tempura lobe (ASM), and centrum semi-ovale of white matter (CSO-WM). The ROIs were registered back into each subject’s space. Then, the number of PVS in those regions were divided by the total ROI volume to extract PVS volume fraction.

**Figure 1.**
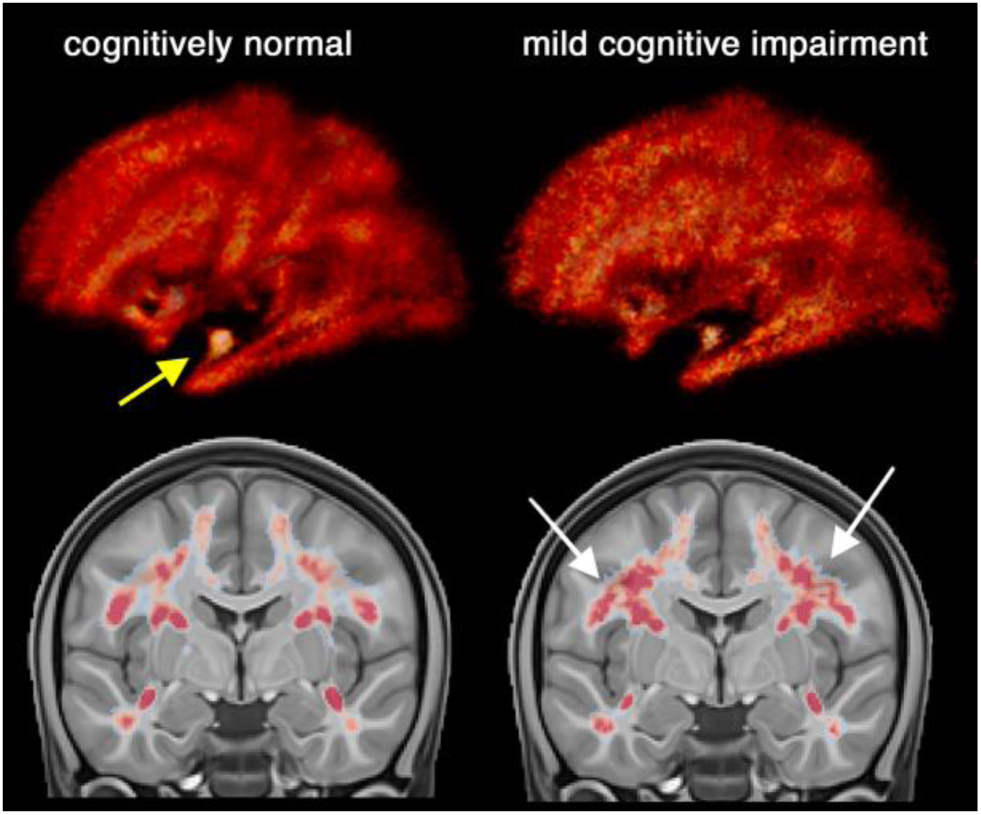
Maps of the perivascular space (PVS) in cognitively normal (CN: n=424) and mild cognitively impaired (MCI: n=173) participants of Alzheimer’s Disease Neuroimaging Initiative 3 (ADNI-3). A higher amount of PVS was observed almost across the entire white matter of the MCI participants and particularly in the centrum semi-ovale region. Surprisingly, in the MCI group, a lower amount of PVS was observed in anterosuperior medial temporal lobe white matter.

All FLAIR and T1w MRI images were manually checked for misalignment and postprocessing failure, randomly and blind to clinical and neuroimaging measures. A total of 37 participants were excluded after the quality control.

We observed MRI manufacturer difference across PVS measures, in which PVS were consistently underestimated in Philips compared to other scanners (*p*<0.01 across all comparisons). Therefore, two post-hoc tests were performed: 1) manufacturer was included in the regression model as covariate. No statistically significant contribution from scanner manufacturer was observed in any statistical model (all *p*-value ≥ 0.55). 2) statistical analysis was repeated, excluding 65 participants that were scanned with the Philips scanner. Statistical conclusions were equivalent to those from the full dataset when images from Philips scanner were excluded (**Supplementary File 3**).

### Statistical analysis

Table 1 summarizes study participants’ demographic and clinical information. To assess the PVS volume fraction differences, we used estimation statistics when comparing CN and MCI group differences [74]. To this end, we estimated the group mean difference by calculating the unpaired Cohen’*d* effect size and the confidence interval between the PVS volume fraction of CN and MCI groups. To calculate the confidence interval of the effect size 5000 bootstrap samples were taken. The confidence interval was bias-corrected and accelerated [74]. We also reported two-sided *p*-value of Mann-Whitney test, which was reported as the likelihood of observing the effect size, if the null hypothesis of zero difference is true. For strong and significant group differences, a secondary statistical analysis was performed to incorporate potential covariates. To this end, we performed logistic regression using cognitive state as dependent variable, adjusting for age, sex, education and brain size. PVS estimates were calculated as a volume fraction, defined as the volume of PVS in the region. For domain-specific analysis, a similar approach with the same covariates was used, but with the difference that CDR was used to make 2 groups: participants with no cognitive deficit (i.e. CDR=0) and participants with mild cognitive deficit (i.e. CDR=0.5).

**Table 1.** Participant summary information.

| Measures | Cognitively Normal | Amnesic MCI |
| --- | --- | --- |
| **N** | 423 | 173 |
| **Female: n (%)** | 252 (59%) | 73 (42%) |
| **Age (years)** | 73.04 ± 7.36 | 75.56 ± 8.16 |
| **Education (years)** | 16.90 ± 2.25 | 15.98 ± 2.78 |
| **Brain volume (mm<sup>3</sup>)** | 1,064,574 ± 107,460 | 1,051,535 ± 106,529 |
| **CDR global\*** | 0.0 (0.0) | 0.5 (0.0) |
| **CDR memory\*** | 0.0 (0.0) | 0.5 (0.0) |
| **CDR sum-of-boxes\*** | 0.0 (0.0) | 1.0 (1.5) |
| **MMSE\*** | 29.0 (1.0) | 29.0 (3.0) |
| **APOE status: n (%)** |  |  |
| ε3/ε3 | 212 (66%) | 78 (61%) |
| ε3/ε4 | 99 (31%) | 37 (29%) |
| ε4/ε4 | 10 (3%) | 12 (9%) |
| **Ab PET (SUVR)<sup>+</sup>** | 1.11 ± 0.17 | 1.16 ± 0.27 |
| **Tau PET (Braak regions 1&2)<sup>++</sup>** | 1.15 ± 0.13 | 1.24 ± 0.21 |
| **Tau PET (Subcortical SUVR)<sup>++</sup>** | 1.13 ± 0.11 | 1.17 ± 0.13 |
| **History of Hypertension: n (%)<sup>+++</sup>** | 91 (34%) | 44 (52%) |
| **Systolic pressure (mm Hg)** | 133.9 ± 17.8 | 134.1 ± 18.0 |
| **Diastolic pressure (mm Hg)** | 75.0 ± 9.6 | 73.4 ± 8.4 |

Values are mean ± SD or frequency (percent).
\* Values are median (IQR) (normality hypothesis was rejected with D'Agostino and Pearson's test [31])
+ n=259 participants had AV45 Aβ PET data: n(CN)=171, n(MCI)=59
++ n=345 participants had AV1451 Tau PET data: n(CN)=240, n(MCI)=79
+++ n=378 participants (269 CN, 86 MCI) had data about history of hypertension
++++ n=508 participants had APOE status data, n=60 with APOE ε2 were excluded in genotype analysis

The association between PVS volume fraction and Tau pathology, as measured by PET, was investigated. For Tau PET analysis, regional SUVRs in medial temporal lobe regions associated with early AD pathology were investigated. In particular, we examined whether PVS volume fraction of ASM is correlated with Tau uptake in regions with early AD pathology: entorhinal cortex (ERC), hippocampus (HC), and parahippocampus (PHC). We also looked at regions associated with Braak stages, as described in the *Positron emission tomography* method section. For these analyses, we formulated a linear regression model to investigate whether PVS volume fraction is a predictor of Flortaucipir (AV-1451) SUVR, adjusting for age, sex, education and brain size. An ordinary least square fitting was used to fit the linear regression to the data using StatsModel Python library. A secondary analysis was also performed including the continuous unthresholded Aβ SUVR as a covariate to compare the effect size of PVS volume fraction against Aβ SUVR.

We then looked into possible explanations for PVS volume fraction mean differences between CN and MCI groups, including amyloid PET, *APOE* status, and cardiovascular measures. For the Amyloid PET analysis, participants were categorized into CN and MCI with and without amyloid presence, which was defined by a cut-off at SUVR of 1.1. Mean PVS volume fractions were then compared across Aβ- and Aβ+ groups, correcting for age, sex, brain volume and education. When the overall association between vascular and PVS volume fraction and amyloid load was assessed, the continuous unthresholded Aβ SUVR values were used. Similar estimation statistics framework was used for *APOE* status comparison and cardiovascular measures. For the *APOE-ε4* carrier status analysis, participants were categorized into three groups based on the number of *APOE ε4* allele repeats: 0 for *ε3/ε3*, 1 for *ε3/ε4*, and 2 for *ε4/ε4*. Participants with *APOE*-*ε2* allele were excluded from genotype analysis as it was shown to be a protective carrier [75, 76]. A secondary analysis was done including *APOE*-*ε2* allele, which resulted to the same conclusion (no association was found).

Finally, because of the pathological changes in blood vessels observed in hypertension as well as the cerebrovascular response to the blood levels of oxygen and carbon dioxide, which are two primary determinants of ventilation [77], we explored the contributions of blood pressure and respiratory rate to the PVS. Available cardiovascular risk factors including history of hypertension and vital signs (systolic and diastolic pressures, respiration rate and temperature) were used as independent variables to explain the variation of PVS volume fraction. Similar linear regression modelling as above was used for this purpose, correcting for age, sex, education and brain volume. The Benjamini–Hochberg procedure with a false discovery rate of 0.05 was used to correct for multiple comparisons (**Supplementary Figure 2** and **Supplementary File 4**). Participants with a history of hypertension had significantly higher PVS volume fraction in the CSO-WM (*p*=0.018, adjusted for age, sex, education and brain volume). After exploring the vital signs (systolic, diastolic and pulse pressure, respiration rate and temperature), we observed a significant inverse association between respiration rate and PVS volume fraction in the ASM (*p*=0.004, adjusted for age, sex, education and brain volume), in which tachypneic participants (respiration rate > 20 per minute) had lower PVS volume fraction in the ASM. After including variables with statistically significant associations with PVS (i.e. history of hypertension and respiration rate) into the analysis of PVS difference by cognitive status (CN vs MCI), all statistical conclusions regarding cognitive group differences remained unchanged.

## Results

A higher total amount of PVS was observed in the normative PVS map of MCI participants compared to CN participants. The increased PVS in MCI compared to CN participants was most evident in the CSO-WM. Surprisingly, PVS was lower in MCI compared to CN in the anterosuperior medial temporal lobe (ASM).

A significantly lower PVS volume fraction in MCI compared to CN groups was observed in both left (*p*=0.00047) and right (*p*=0.0009) anterosuperior medial temporal lobe (corrected for age, sex, education and brain volume). We also observed an asymmetrical pattern of PVS volume fraction in this region in both CN (*p*=7e-13) and MCI (*p*=0.0001) groups (higher on the right hemisphere). No asymmetry was observed in the CSO-WM (*p*=0.6). PVS volume fraction of the CSO-WM was higher in MCI compared to CN (**Figure 2.d**), but not statistically significant after adjusting for age, sex, education and brain volume (*p*=0.07). When participants were categorized based on CDR scores (**Figure 2.f and 2.g**), the same pattern was observed: PVS volume fraction of ASM were significantly lower in participants with signs of cognitive deficit (CDR=0.5) compared with cognitively normal individuals (CDR=0) (left ASM: *p*=0.00014, right ASM: *p*=0.00025, corrected for age, sex, education and brain volume). No significant differences were observed in the CSO-WM after adjusting for aforementioned covariates (**Figure 2.h**).

**Figure 2.**
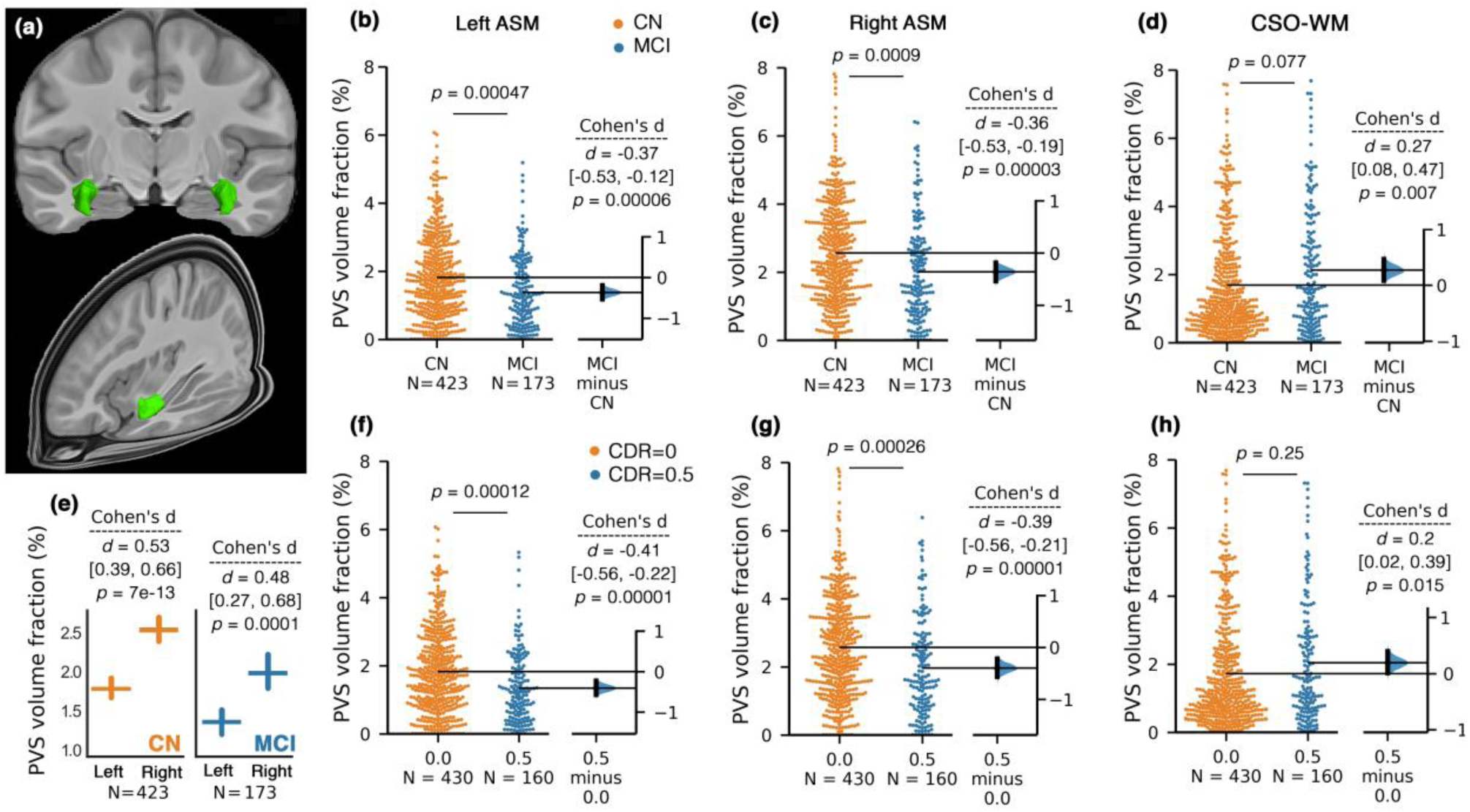
PVS volume fraction in anterosuperior medial temporal lobe (ASM) was significantly lower in mild cognitive impairment (MCI) participants compared to cognitively normal (CN). A 3D render of the ASM on template space is presented in **(a)**. Bilateral and significant PVS volume fraction differences between CN and MCI participants were observed in ASM **(b** and **c)**. Unpaired Cohen’s *d* effect size and confident intervals were reported (*d* [CI]) for the group differences between CN and MCI using estimation statistics [74]. Two-sided *p*-value of Mann-Whitney test was also reported. The second *p*-value (reported on the line between the swarm plots) is adjusted for age, sex, years of education and brain size using logistic regression. The PVS volume fraction in ASM was asymmetrical across both CN and MCI groups **(e)**; mean and standard errors are shown. Diagnostic group comparison on PVS volume fraction across the centrum semi-ovale of the white matter (CSO-WM) is shown in **(d)**. **(f-h)** plots compare PVS volume fraction of the ASM when study participants were categorized based on clinical dementia rating (CDR).

PVS volume fraction in the ASM was significantly associated with Tau PET standardized uptake value ratio (SUVR) in the entorhinal cortex (ERC), which is a pathological hallmark of AD in the first stages (Braak stages I and II). The association appeared to be lateralized, in which only the right ERC showed significant correlation (beta (SE) = -0.02 (0.007); *p*=0.006, corrected for age, sex, education and brain volume). Unlike ERC, we did not observe significant associations between PVS volume fraction and Tau uptake in the hippocampus and parahippocampal cortex nor in the white matter. ASM PVS volume fraction was associated with Tau uptake in regions associated with Braak I and II (i.e. HC and ERC combined), which is likely dominated by the significant association of PVS volume fraction and the ERC Tau uptake. The correlation was weaker for Braak stage III and IV, which includes parahippocampal, fusiform, and lingual gyri as well as the amygdala. The weakest correlation was found in regions affected in the latest Braak stages (V and VI), suggesting that ASM PVS alteration could be a biomarker of early AD pathology (the list of the brain regions analyzed according to the Braak stages are included in **Supplementary File 2**).

PVS volume fractions across different regions were independent of Amyloid load or *APOE* ε4 status (**Figure 4** and **5**). A secondary linear regression was performed using cerebral amyloid load values (Aβ SUVR) as a predictor of PVS volume fraction, which showed that Aβ uptake was not a predictor of PVS volume fraction (*p*=0.61). The same conclusion was drawn when the analysis was performed on CN and MCI groups separately. No differences in mean PVS volume fraction between Aβ- and Aβ+ participants were observed in MCI and CN participants (CN (left ASM): *p*=0.75, CN (right ASM): *p*=0.98, MCI (left ASM): *p*=0.55, MCI (right ASM): *p*=0.16, CN (CSO-WM): *p*=0.95, MCI (CSO-WM): *p*=0.15, **Supplementary Figure 3**). We also observed no PVS volume fraction difference across participants with different *APOE* e4 allele repeats (**Figure 5**).

**Figure 3.**
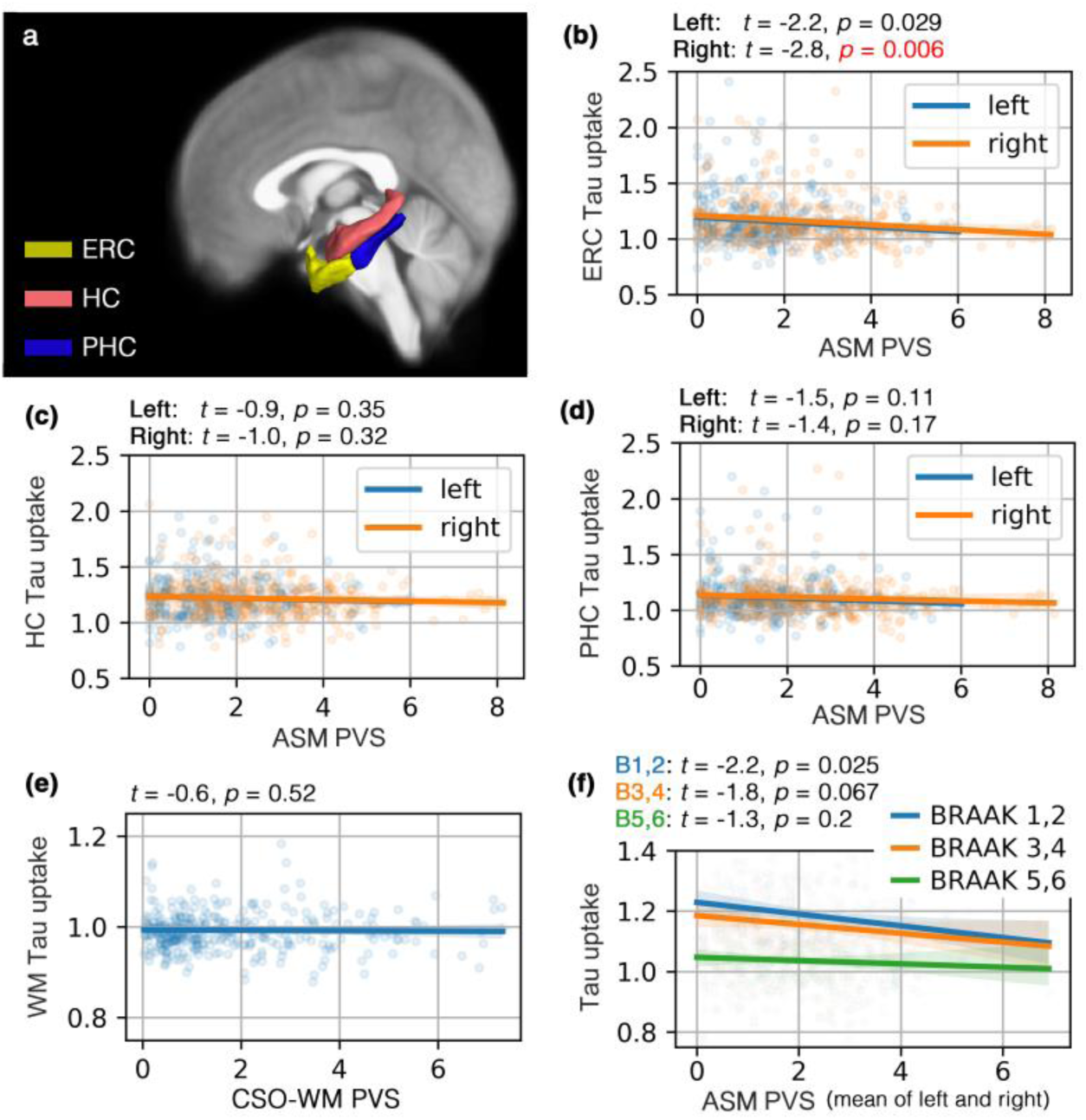
PVS volume fraction was associated with early AD pathology. Studied regions of interest are shown in **(a)**. PVS volume fraction of the anterosuperior medial temporal lobe (ASM) was a predictor of Tau uptake (AV1451 PET Tau SUVR) in entorhinal cortex (ERC) **(b)** (right ASM: *p*=0.006, left ASM: *p*=0.029, corrected for age, sex, education and brain size), in contrast with PVS volume fraction of the centrum semi-ovale of the white matter (CSO-WM) (*p*=0.52) **(e)**. A non-significant correlation between PVS volume fraction of ASM and Tau uptake in hippocampus (HC) and parahippocampal cortex (PHC) was observed **(c-d)**. Participants with lower PVS volume fraction in ASM appeared to have higher amount of Tau in ERC. The correlation between PVS volume fraction and Tau SUVR was highest in regions involved in early AD pathology (i.e. ERC as per Braak stage I), and the association was weaker for the regions affected in later stages (Braak stage V and VI), characterized by extensive neocortical involvement **(f)**. The association analysis was performed in a subset of the population including participants with both Tau PET and MRI data (n=319; n(CN)=240, n(MCI)=79).

**Figure 4.**
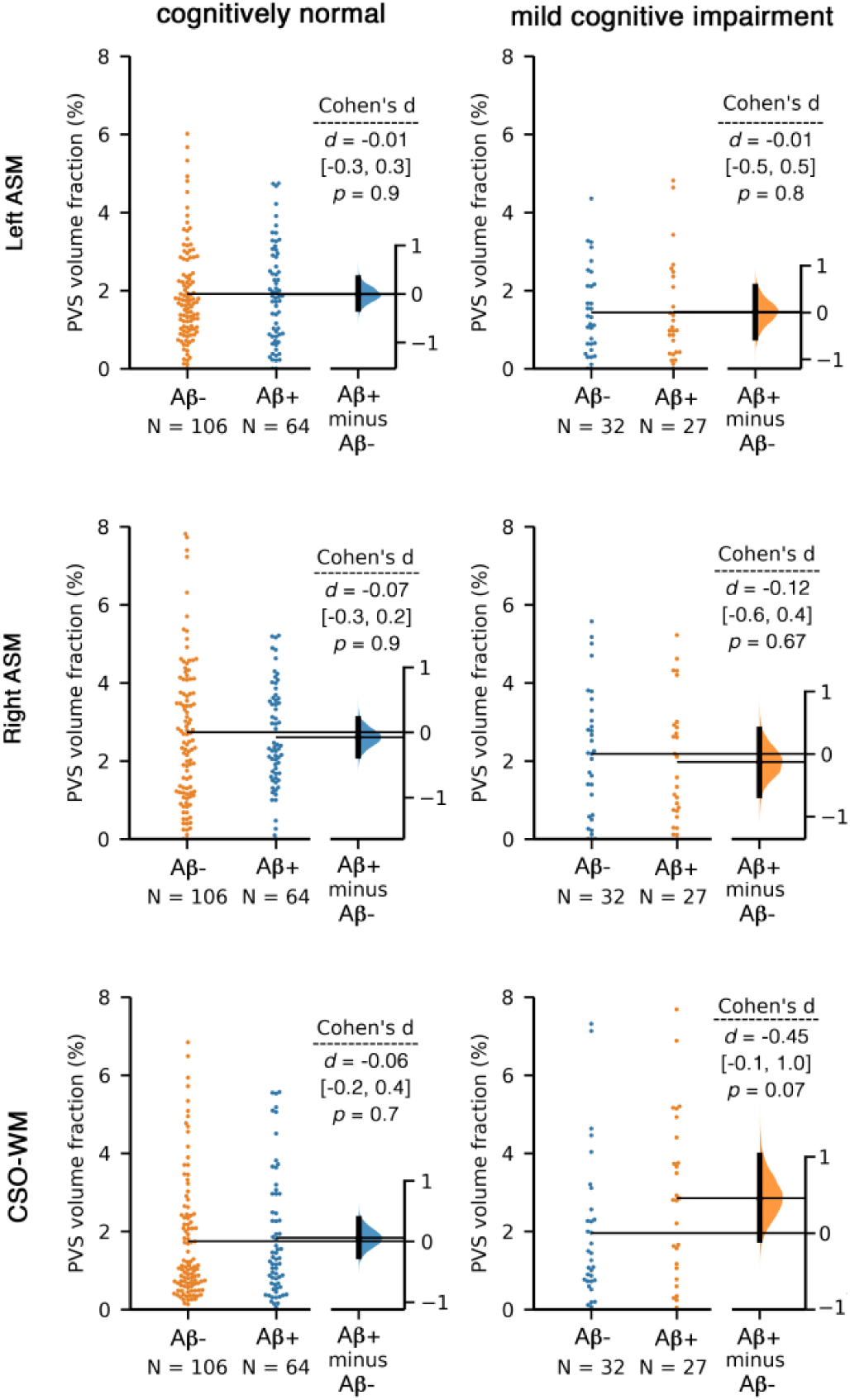
PVS volume fraction of the anterosuperior medial temporal lobe (ASM) and centrum semi-ovale of the white matter (CSO-WM) were not associated with the amyloid beta uptake, as measured by positron emission tomography. Aβ-/+ categorization was applied based on Florbetapir cortical summary measurement (SUVR). Florbetapir cutoff of 1.11 using the whole cerebellum region as a reference. Equivalent statistical conclusions were drawn when we used a cutoff of 0.79 in a composite reference region including cerebellum, brainstem/pons, and eroded subcortical white matter. Linear regression analysis using Aβ SUVR also showed no association between Aβ uptake and PVS volume fraction (*p*=0.61). When regional values of Aβ SUVR were used (instead of global SUVR), no associations were observed (*p*>0.6 for all tests).

**Figure 5.**
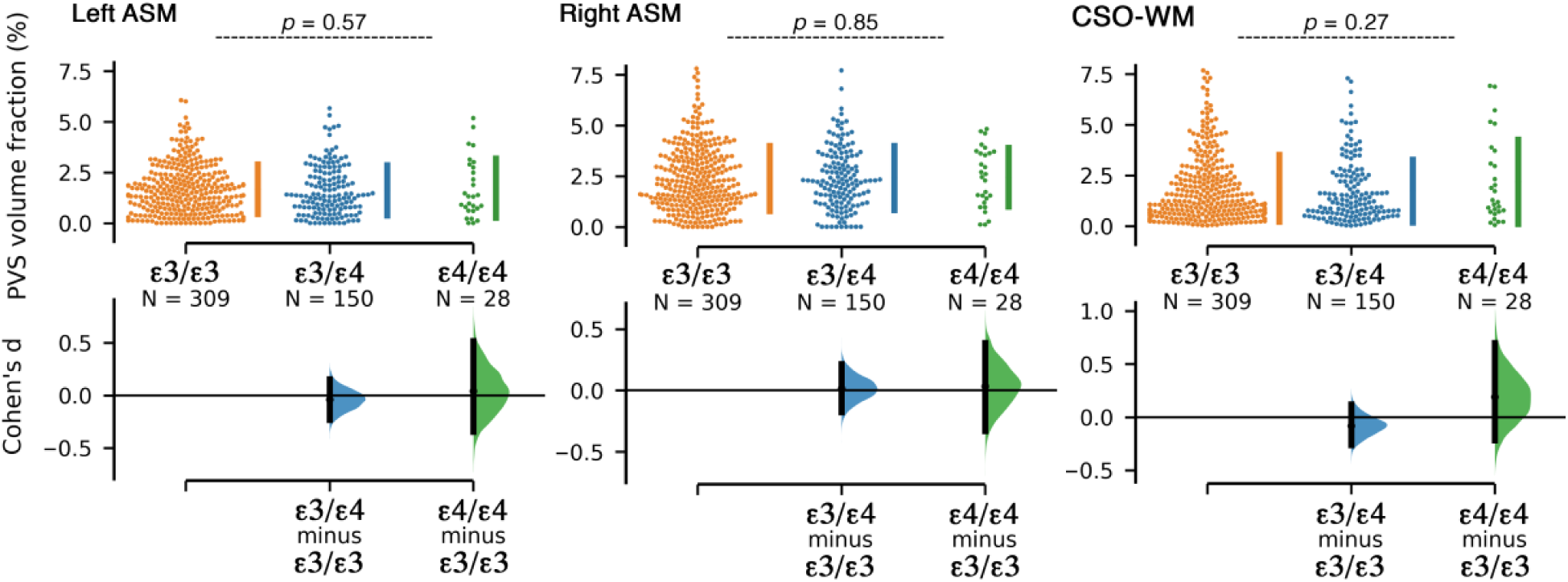
PVS volume fraction of the anterosuperior medial temporal lobe (ASM) and centrum semi-ovale of the white matter (CSO-WM) were not associated with *APOE* ε4 allele repeats. Participants were categorized based on the number of *APOE* ε4 allele repeats.

Finally, when total ASM PVS volume fraction and composite cerebral amyloid load values (Aβ SUVR) were included in the same logistic regression model to predict the cognitive status (MCI vs CN), the ASM PVS (beta (SE) = -0.33 (0.1); *t*=-2.3; *p*=0.019; lower in MCI) was a stronger predictor than that of PET SUVR (beta (SE) = 2.26 (1.2); *t*=1.8; *p*=0.07; higher in MCI), supporting the hypothesis that alterations in the supply and clearance of the ASM could be an early biomarker of AD-related cognitive decline. No association between PVS volume fraction and CSF biomarkers were found (**Supplementary File 5**). Continuous data of CSF Aβ_1-42_, Aβ_1-40_, Tau and phosphorylated Tau (p-Tau) were used for the association analysis, adjusting for age, sex, years of education and brain size.

## Discussion

In summary, our findings show that older adults with mild cognitive impairment develop alterations in the PVS; specifically, a significantly lower PVS volume fraction was found in ASM and a (non-significantly) higher PVS volume fraction was measured in the CSO-WM compared with cognitively normal participants. Since group differences in ASM PVS measurements between CN and MCI were significant after correcting for several covariates, including age and brain volumes, these data indicate that PVS alterations in ASM reflect cognitive impairment independent of normal aging and may therefore represent a sensitive biomarker of early cognitive dysfunction. Concerning the PVS in CSO-WM, our present results are consistent with previous studies showing an association between enlarged PVS in CSO-WM and both Alzheimer’s disease and cerebral amyloid angiopathy [26, 78]; however, the enlargement of PVS in CSO-WM of MCI individuals in our cohort was not significant after correction for brain volume, suggesting that these changes might actually be a consequence related to the atrophy of the brain, as previously postulated [79, 80]. The CSO-WM results also invite further region-specific analysis across the white matter as neuroimaging techniques become more sensitive to PVS presence [68]. Intriguingly, for the first time we demonstrated a loss of PVS signal in the medial temporal lobe, a critical region intimately connected with the limbic system and anatomically in contact with the hippocampus. The PVS volume fraction of this region was a significant predictor of the cognitive status in a multivariate logistic regression model. Since the signal measured in PVS strictly depends on the CSF signal, we hypothesize that the reduction in PVS volume fraction may represent a consequence of an occlusion and/or obstruction of the perivascular spaces in ASM, resulting in impaired CSF flow and loss of CSF signal. Montagne et al. have previously demonstrated that BBB permeability as measured by MRI is significantly increased in the hippocampus of pericyte-deficient mice, resulting in the early accumulation of blood-derived toxic fibrin(ogen) deposits in the cerebral parenchyma and perivascular spaces, which ultimately leads to diminished blood flow and white matter damage [1]. Although it is not known whether an early BBB breakdown similarly occurs in ASM, we believe that the obstruction of the PVS by these deposits may impair the CSF perivascular flow, with resulting decreased detectability of PVS in ASM of MCI individuals. In addition, we found that these PVS changes were independent from the Aβ and Tau pathology, as measured by PET and CSF. This is consistent with the recent demonstration that neurovascular dysfunction and BBB breakdown occur early in AD and independently of Aβ and Tau [2], and therefore supports our pathogenic hypothesis. Interestingly, only the Tau uptake in the entorhinal cortex, a region classically described to be affected early according to the Braak staging, showed a significant association with the PVS changes in ASM, suggesting that the PVS alteration in ASM may represent an early biomarker of AD pathology. It has also been proposed that the impairment of the glymphatic flow through the PVS might reduce the clearance of metabolic waste products, including Aβ and Tau [5], whose deposits could further impair the perivascular drainage, leading to ulterior accumulation in a ‘feed-forward’ process. However, whether BBB breakdown and perivascular flow impairment with reduced clearance are related and to which extent these two factors contribute to neurodegeneration remain to be explored by future studies.

According to our results the *APOE* haplotype did not influence the PVS volume fraction and distribution in ASM and CSO-WM. Pathological changes in the structure and/or function of cerebral microvasculature has been shown to play a major role in dementia [21,81,82]. PVS, as direct biomarker of the cerebral glia-lymphatic pathway and indirect biomarker of cerebrovascular health, represent a “bridge” between two key mechanisms underlying neurodegeneration and may therefore be a novel indicator of early pathophysiological processes leading to cognitive impairment. On one hand, BBB breakdown in medial temporal regions has been demonstrated to affect a number of cognitive functions, such as working memory, executive function, and semantic fluency tasks, as a consequence of disrupted structures and connecting pathways [2]; BBB damage may indeed induce neurotoxicity and axonal degeneration via multiple mechanisms, including perivascular and parenchymal leakage of blood-borne neurotoxic proteins as well as activation of inflammatory responses [6]. For example, plasmin and fibrinogen derived from the circulating blood are able to provoke neuronal injury, axonal retraction, and further BBB damage [83, 84]; moreover, influx of albumin leads to perivascular edema, which obstructs not only the perivascular flow but also the microcirculation blood flow, resulting in hypoxia and neuronal death [28, 85].

On the other hand, the impairment of the perivascular drainage system mediating waste products elimination and Aβ clearance is responsible for cerebral amyloid angiopathy, which is considered a pathological hallmark of AD and is related to cognitive decline [86–88]. Since in both cases pathological changes in PVS (i.e., obstruction/occlusion and/or enlargement), are reasonably expected as a consequence of the accumulation of proteins and other substances in the perivascular compartment, it is important to analyze and quantify these alterations that could be used not only as a potential biomarker of cognitive decline, but also for a better understanding of the pathophysiology and consequently for planning appropriate therapeutic interventions.

Future longitudinal studies, focusing on examining the alteration of the PVS as a function of pathophysiology and the cognitive decline, are demanded to further validate our findings and to evaluate the efficacy of PVS mapping as a surrogate for brain clearance and vascular health. The precise quantification and mapping of PVS offer the possibility to significantly improve the characterization of PVS changes, which were previously underappreciated by the visual rating methods. Moreover, the development of automated pipelines for computing unbiased and reproducible measurements of PVS will facilitate the analysis of PVS in multi-center studies and the comparison of the results with those obtained by other research groups, leading to more solid and comprehensive knowledge of the role of PVS. However, for this purpose it is fundamental to consider that the MRI-based PVS analysis strongly relies on the image resolution: higher resolution corresponds to increased PVS detection and improved segmentation [68,90,91]. Small PVS remain still undetected on MRI or are not discernible from noise. This limitation currently does not allow to identify the submillimeter pathological changes from clinical data, which are hypothetically supposed to occur at the very early pre-clinical stages of the disease. Finally, quantitative characteristics of the PVS using non-invasive techniques, such as diffusion MRI, could be used to extract submillimeter information [91, 92].

## Conclusions

In conclusion, our results demonstrate that PVS may represent an important new biomarker of cognitive decline, in individuals in the Alzheimer’s continuum as well as those without Aβ and/or pTau positivity. In contrast with CSF and neuroimaging biomarkers based on contrast-enhanced MRI, this novel biomarker is completely non-invasive and does not require any contrast injection since it can be accurately measured using conventional clinical sequences. This discovery encourages further investigation of the cerebral PVS in terms of both pathophysiology and new potential therapeutic strategies for cognitive dysfunction.

## Acknowledgement

This work was supported by NIH grants: 2P41EB015922-21, 1P01AG052350-01 and USC ADRC 5P50AG005142. The content is solely the responsibility of the authors and does not necessarily represent the official views of the NIH.

**ADNI:** Data collection and sharing for this project was funded by the Alzheimer’s Disease Neuroimaging Initiative (ADNI) (National Institutes of Health Grant U01 AG024904) and DOD ADNI (Department of Defense award number W81XWH-12-2-0012). ADNI is funded by the National Institute on Aging, the National Institute of Biomedical Imaging and Bioengineering, and through generous contributions from the following: AbbVie, Alzheimer’s Association; Alzheimer’s Drug Discovery Foundation; Araclon Biotech; BioClinica, Inc.; Biogen; Bristol-Myers Squibb Company; CereSpir, Inc.; Cogstate; Eisai Inc.; Elan Pharmaceuticals, Inc.; Eli Lilly and Company; EuroImmun; F. Hoffmann-La Roche Ltd and its affiliated company Genentech, Inc.; Fujirebio; GE Healthcare; IXICO Ltd.; Janssen Alzheimer Immunotherapy Research & Development, LLC.; Johnson & Johnson Pharmaceutical Research & Development LLC.; Lumosity; Lundbeck; Merck & Co., Inc.; Meso Scale Diagnostics, LLC.; NeuroRx Research; Neurotrack Technologies; Novartis Pharmaceuticals Corporation; Pfizer Inc.; Piramal Imaging; Servier; Takeda Pharmaceutical Company; and Transition Therapeutics. The Canadian Institutes of Health Research is providing funds to support ADNI clinical sites in Canada. Private sector contributions are facilitated by the Foundation for the National Institutes of Health (www.fnih.org). The grantee organization is the Northern California Institute for Research and Education, and the study is coordinated by the Alzheimer’s Therapeutic Research Institute at the University of Southern California. ADNI data are disseminated by the Laboratory for Neuro Imaging at the University of Southern California.

---

**Supplementary Figure 1.**
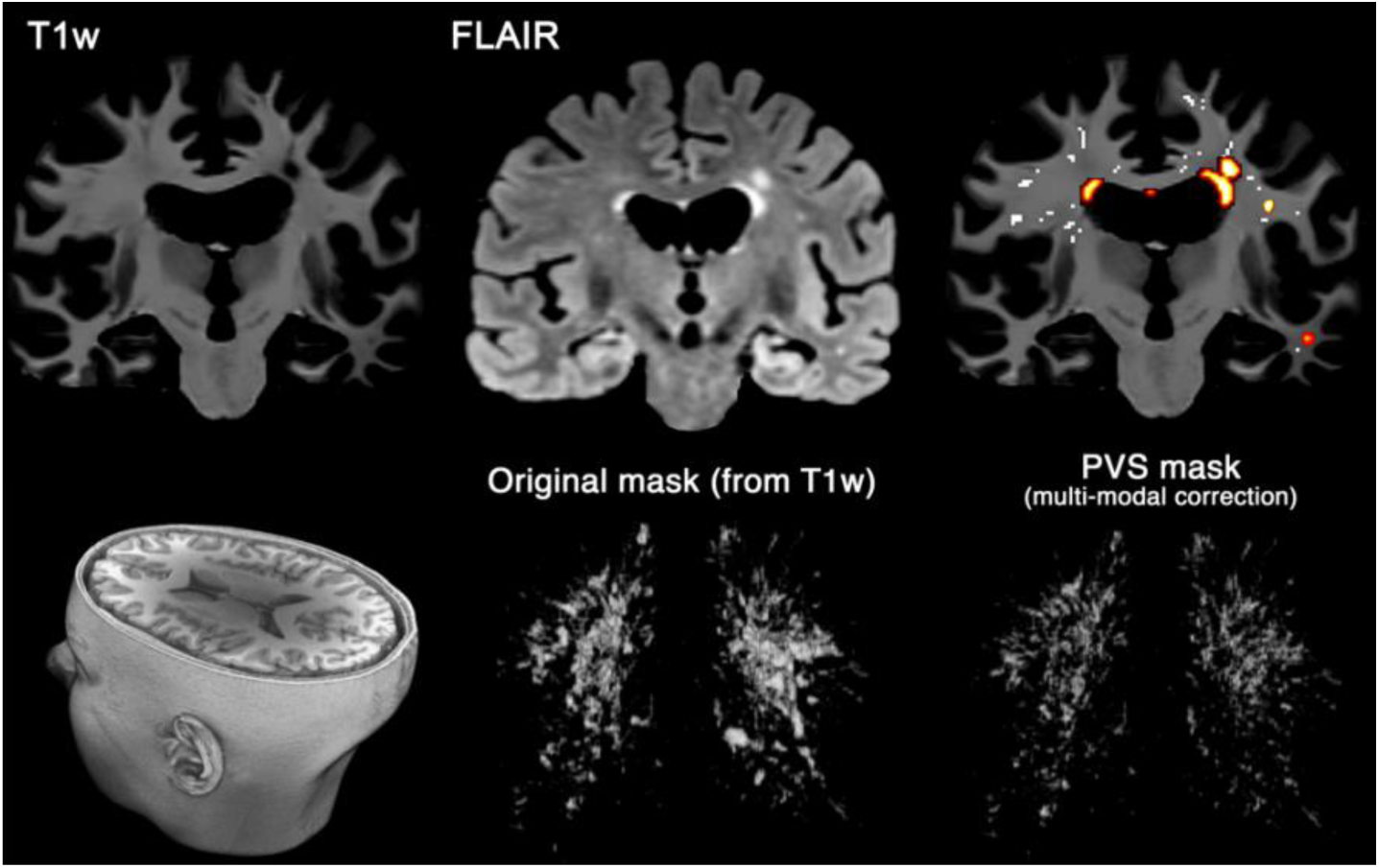
Multi-modal PVS segmentation of clinical data. Perivascular spaces (PVS) were mapped from T1w images and then white matter hyperintensities, which was falsely segmented as PVS, were excluded using FLAIR images. PVS segmentation was done using an automated and highly reliable quantification technique that we developed and validated recently (38), based on non-local filtering, Frangi filtering and optimized mask identification. For each segmented voxel in T1w, corresponding FLAIR voxels was checked, and if the FLAIR voxel value was a white matter hyperintensity, then the voxel was excluded from the final PVS mask.

**Supplementary Figure 2.**
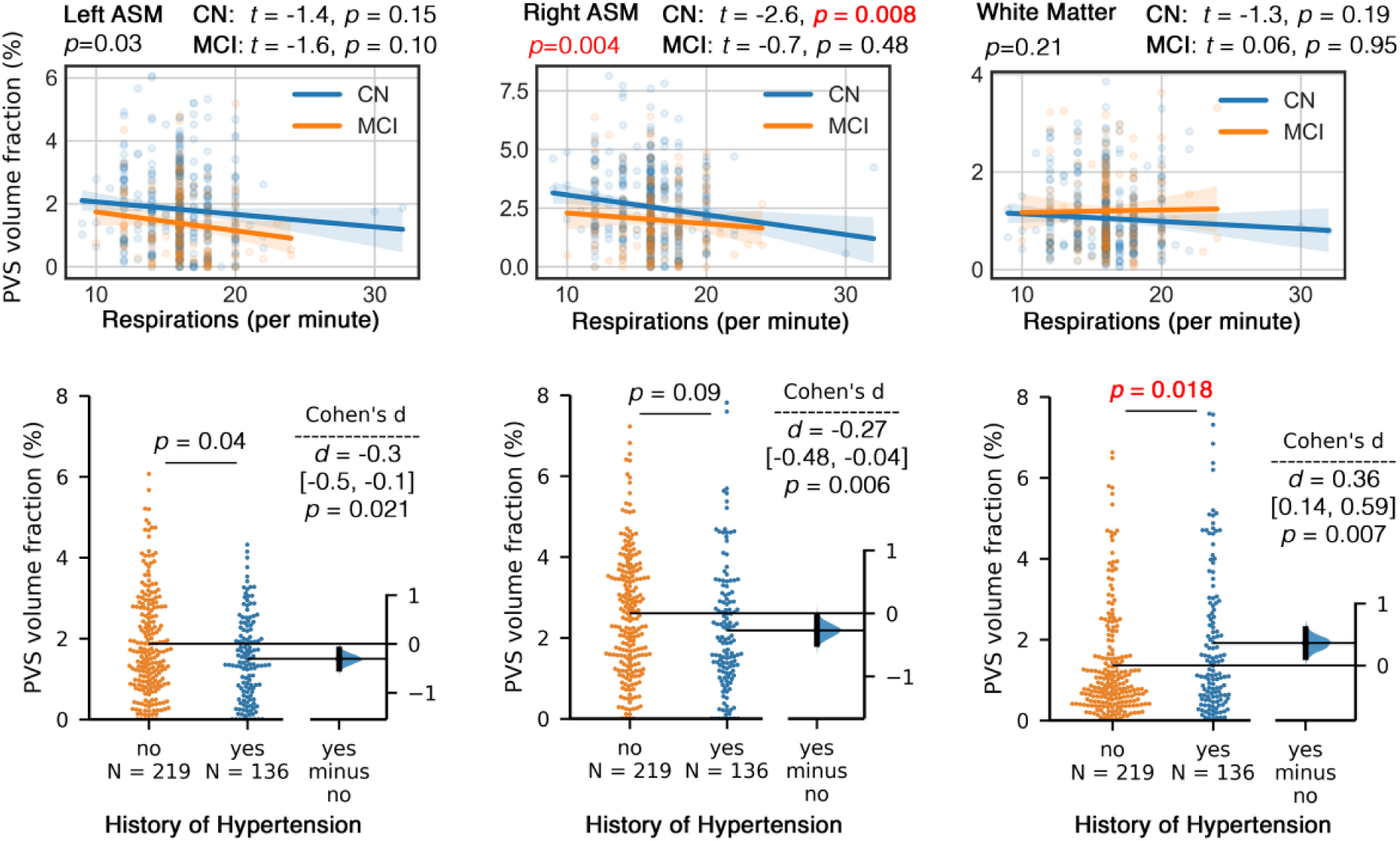
History of hypertension and respiration rate are differentially associated with the PVS volume fraction. **First row**: respiratory rate (number of breaths per minute) is negatively associated with PVS volume fraction in anterosuperior medial temporal lobe (ASM), but not associated with PVS volume fraction in centrum semi-ovale of the white matter. **Second row**: Subjects with history of hypertension had significantly higher PVS volume fraction in centrum semi-ovale of white matter, but PVS volume fraction in ASM did not differ by hypertension (not significant after multi-comparison correction). Statistics that survived the multiple-comparison correction using Benjamini–Hochberg procedure were colorcoded red.

**Supplementary Figure 3.**
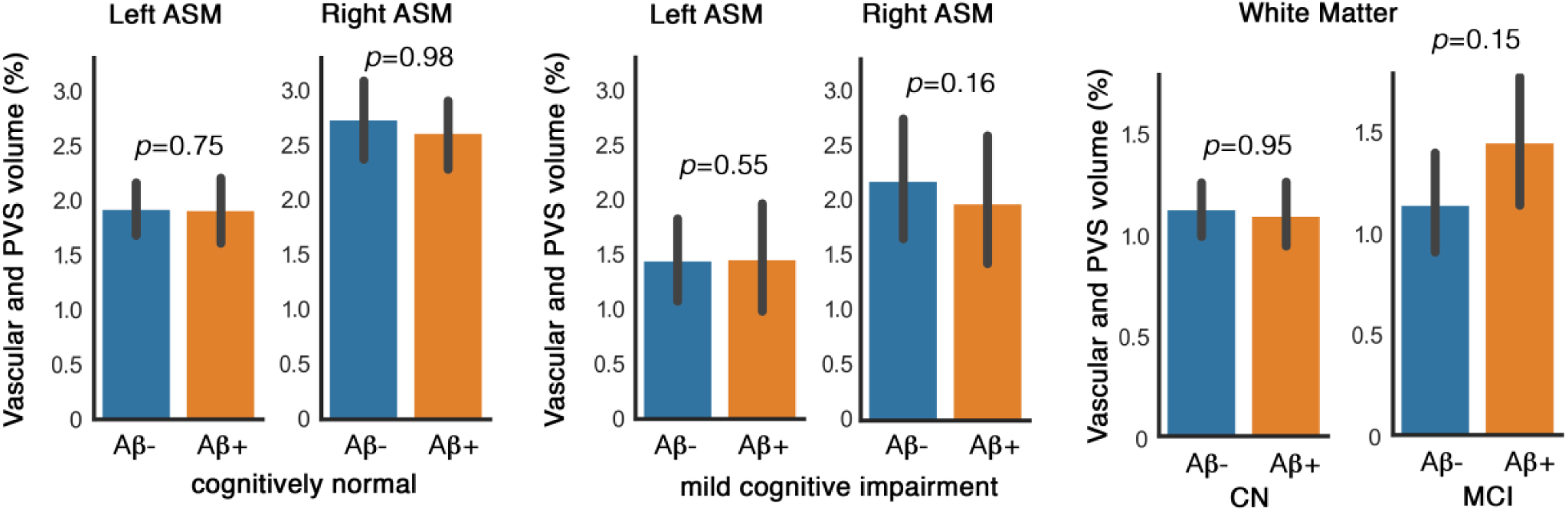
PVS volume fraction of Aβ- and Aβ+ subjects. Vascular and PVS volume fraction of the anterosuperior medial temporal lobe (ASM) and centrum semi-ovale of the white matter were not associated with the amyloid beta update, as measured by positron emission tomography.

**Supplementary File 1.** Inclusion/Exclusion Criteria:

Men and women across CN, amnestic MCI, and AD participant groups were included. Inclusion criteria were:

1. Geriatric Depression Scale score less than 6.
2. Age between 55-90 years (inclusive).
3. Study partner who has frequent contact with the participant (i.e., minimum average of 10 hours per week) and is available to accompany the participant to all clinic visits for the duration of the protocol.
4. Visual and auditory acuity adequate for neuropsychological testing.
5. Good general health with no diseases expected to interfere with the study.
6. Participant is not pregnant, lactating, or of childbearing potential (i.e. women must be two years post-menopausal or surgically sterile).
7. Willing and able to participate in a longitudinal imaging study.
8. Modified Hachinski Ischemic Score less than or equal to 4.
9. Completed six grades of education or has a good work history (sufficient to exclude mental retardation).
10. Must speak English or Spanish fluently.
11. Willing to undergo repeated MRIs (3Tesla) and at least two PET scans
12. Agrees to collection of blood for genomic analysis (including GWAS sequencing and other analysis), APOE testing and biospecimen banking.
13. Agrees to collection of blood for biomarker testing.
14. Agrees to at least one lumbar puncture for the collection of CSF.
15. Agrees to share genomic data and biomarker samples.

The following additional exclusion criteria apply to all diagnostic categories:

1. Screening/Baseline MRI brain scan with evidence of infection, infarction, or other focal lesions or multiple lacunes or lacunes in a critical memory structure
2. Subjects that have any contraindications for MRI studies, including the presence of cardiac pacemakers, or metal fragments or foreign objects in the eyes, skin or body
3. Major depression, bipolar disorder as described in DSM-IV within the past 1 year. Psychotic features, agitation or behavioral problems within the last 3 months that could lead to difficulty complying with the protocol.
4. Currently treated with medication for obsessive-compulsive disorder or attention deficit disorder.
5. History of schizophrenia (DSM IV criteria).
6. History of alcohol or substance abuse or dependence within the past 2 years (DSM IV criteria).
7. Any significant systemic illness or unstable medical condition, which could lead to difficulty complying with the protocol.
8. Clinically significant abnormalities in B12 or thyroid function tests that might interfere with the study. A low B12 is exclusionary, unless follow-up labs (homocysteine and methylmalonic acid) indicate that it is not physiologically significant.
9. Residence in a skilled nursing facility.
10. Current use of specific psychoactive medications (e.g., certain antidepressants, neuroleptics, chronic anxiolytics or sedative hypnotics). Current use of warfarin or other anticoagulants such as dabigatran, rivaroxaban and apixaban (exclusionary for lumbar puncture).
11. Current use of any other exclusionary medications
12. Investigational agents are prohibited one month prior to entry and for the duration of the trial.
13. Participation in clinical studies involving neuropsychological measures being collected more than one time per year.

**Supplementary File 2**. FreeSurfer-defined region codes for Braak regions of interest:

**Braak 1 and 2 composite region (Braak12):**

**Braak 1**

1006 L_entorhinal

2006 R_entorhinal

**Braak 2**

17 L_hippocampus

53 R_hippocampus

**Braak 3 and 4 composite region (Braak34):**

**Braak 3**

1016 L_parahippocampal

1007 L_fusiform

1013 L_lingual

18 L_amygdala

2016 R_parahippocampal

2007 R_fusiform

2013 R_lingual

54 R_amygdala

**Braak 4**

1015 L_middletemporal

1002 L_caudantcing

1026 L_rostantcing

1023 L_postcing

1010 L_isthmuscing

1035 L_insula

1009 L_inferiortemporal

1033 L_temppole

2015 R_middletemporal

2002 R_caudantcing

2026 R_rostantcing

2023 R_postcing

2010 R_isthmuscing

2035 R_insula

2009 R_inferiortemporal

2033 R_temppole

**Braak 5 and 6 composite region (Braak56):**

**Braak 5**

1028 L_superior_frontal

1012 L_lateral_orbitofrontal

1014 L_medial_orbitofrontal

1032 L_frontal_pole

1003 L_caudal_middle_frontal

1027 L_rostral_middle_frontal

1018 L_pars_opercularis

1019 L_pars_orbitalis

1020 L_pars_triangularis

1011 L_lateraloccipital

1031 L_parietalsupramarginal

1008 L_parietalinferior

1030 L_superiortemporal

1029 L_parietalsuperior

1025 L_precuneus

1001 L_bankSuperiorTemporalSulcus

1034 L_tranvtemp

2028 R_superior_frontal

2012 R_lateral_orbitofrontal

2014 R_medial_orbitofrontal

2032 R_frontal_pole

2003 R_caudal_middle_frontal

2027 R_rostral_middle_frontal

2018 R_pars_opercularis

2019 R_pars_orbitalis

2020 R_pars_triangularis

2011 R_lateraloccipital

2031 R_parietalsupramarginal

2008 R_parietalinferior

2030 R_superiortemporal

2029 R_parietalsuperior

2025 R_precuneus

2001 R_bankSuperiorTemporalSulcus

2034 R_tranvtemp

**Braak 6**

1021 L_pericalcarine

1022 L_postcentral

1005 L_cuneus

1024 L_precentral

1017 L_paracentral

2021 R_pericalcarine

2022 R_postcentral

2005 R_cuneus

2024 R_precentral

2017 R_paracentral

**Supplementary File 3.** Statistical analysis results of when PVS volume fraction of ASM were used to predict cognitive state, using a logistic regression, using age, sex, education and brain volume as covariates - excluding images acquired with Philips scanner.

|  | **coef** | **Std err** | **z** | **P-value** | **0.025 CI** | **0.975 CI** |
| --- | --- | --- | --- | --- | --- | --- |
| **Left ASM** | -0.34 | 0.093 | -3.738 | 0.000186 | -0.529 | -0.165 |
| **Right ASM** | -0.2045 | 0.070 | -2.908 | 0.004 | -0.342 | -0.067 |
| **CSO-WM** | 0.109 | 0.059 | 1.867 | 0.062 | -0.005 | 0.225 |

**Supplementary File 4.** Statistical analysis results of the association between PVS volume fraction and health measures (e.g. vital signs), with age, sex, education and brain volume as covariates. Here we tested if below test measures are predictors of PVS volume fraction.

**Systolic pressure (N= 592)**
|  | **coef** | **Std err** | **z** | **P-value** | **0.025 CI** | **0.975 CI** |
| --- | --- | --- | --- | --- | --- | --- |
| **Left ASM** | -0.0030 | 0.003 | -1.017 | 0.309 | -0.009 | 0.003 |
| **Right ASM** | 0.0002 | 0.004 | 0.055 | 0.956 | -0.007 | 0.008 |
| **CSO-WM** | 0.0035 | 0.004 | 0.898 | 0.369 | -0.004 | 0.011 |

**Diastolic pressure (N= 592)**
|  | **coef** | **Std err** | **z** | **P-value** | **0.025 CI** | **0.975 CI** |
| --- | --- | --- | --- | --- | --- | --- |
| **Left ASM** | -0.0004 | 0.005 | -0.073 | 0.942 | -0.011 | 0.010 |
| **Right ASM** | 0.0030 | 0.007 | 0.445 | 0.657 | -0.010 | 0.016 |
| **CSO-WM** | -0.0072 | 0.007 | -1.016 | 0.310 | -0.021 | 0.007 |

**Pulse pressure (N= 592)**
|  | **coef** | **Std err** | **z** | **P-value** | **0.025 CI** | **0.975 CI** |
| --- | --- | --- | --- | --- | --- | --- |
| **Left ASM** | -0.0038 | 0.003 | -1.124 | 0.261 | -0.010 | 0.003 |
| **Right ASM** | -0.0009 | 0.004 | -0.217 | 0.828 | -0.009 | 0.007 |
| **CSO-WM** | 0.0075 | 0.004 | 1.677 | 0.094 | -0.001 | 0.016 |

**Temperature (N=587)**
|  | **coef** | **Std err** | **z** | **P-value** | **0.025 CI** | **0.975 CI** |
| --- | --- | --- | --- | --- | --- | --- |
| **Left ASM** | 0.1686 | 0.119 | 1.419 | 0.156 | -0.065 | 0.402 |
| **Right ASM** | -0.0903 | 0.150 | -0.600 | 0.548 | -0.386 | 0.205 |
| **CSO-WM** | 0.4176 | 0.158 | 2.639 | 0.009 | 0.107 | 0.729 |

**Supplementary File 5.**
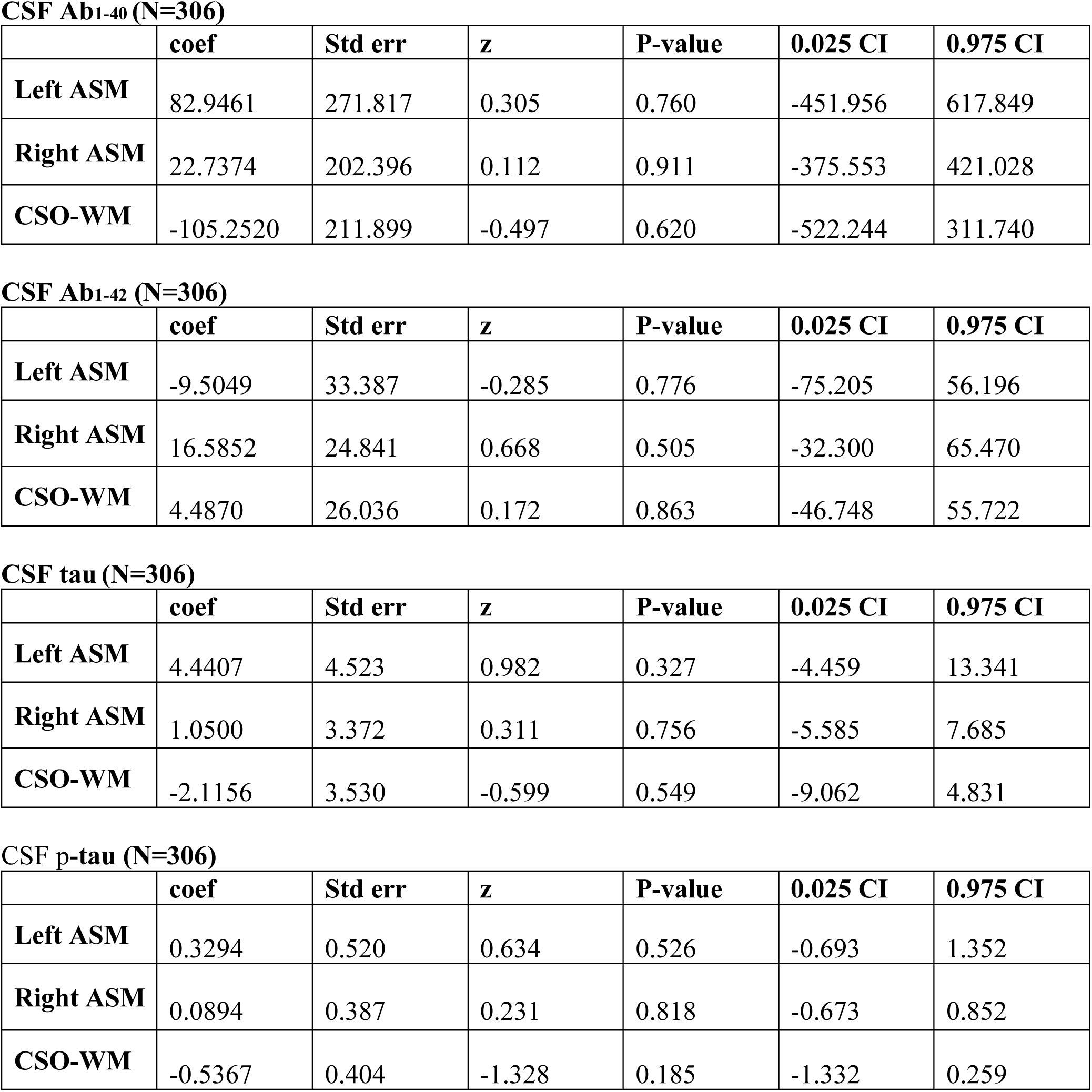
Statistical analysis results of the association between PVS volume fraction and CSF biomarkers, using age, sex, education and brain volume as covariates.

